# The use of non-functional clonotypes as a natural calibrator for quantitative bias correction in adaptive immune receptor repertoire profiling

**DOI:** 10.1101/2021.03.24.436794

**Authors:** Anastasia Smirnova, Anna Miroshnichenkova, Yulia Olshanskaya, Michael Maschan, Yuri Lebedev, Dmitriy Chudakov, Ilgar Mamedov, Alexander Komkov

## Abstract

High-throughput sequencing of adaptive immune receptor repertoires is a valuable tool for receiving insights in adaptive immunity studies. Several powerful TCR/BCR repertoire reconstruction and analysis methods have been developed in the past decade. However, detecting and correcting the discrepancy between real and experimentally observed lymphocyte clone frequencies is still challenging. Here we discovered a hallmark anomaly in the ratio between read count and clone count-based frequencies of non-functional clonotypes in multiplex PCR-based immune repertoires. Calculating this anomaly, we formulated a quantitative measure of V- and J-genes frequency bias driven by multiplex PCR during library preparation called Over Amplification Rate (OAR). Based on the OAR concept, we developed an original software for multiplex PCR-specific bias evaluation and correction named iROAR: Immune Repertoire Over Amplification Removal (https://github.com/smiranast/iROAR). The iROAR algorithm was successfully tested on previously published TCR repertoires obtained using both 5’ RACE (Rapid Amplification of cDNA Ends)-based and multiplex PCR-based approaches and compared with a biological spike-in-based method for PCR bias evaluation. The developed approach can increase the accuracy and consistency of repertoires reconstructed by different methods making them more applicable for comparative analysis.

## Introduction

Adaptive immune receptor (TCR – T-cell receptor and BCR – B-cell receptor) repertoire is usually defined as a set of TCR or BCR sequences obtained from an individual’s blood, bone marrow, or specific lymphocyte population. Reflecting the T/B cell’s clonal composition, the repertoire is characterized by a high degree of specificity for each individual and substantial variation in clone frequencies. The accuracy of both sequences and frequencies of TCR/BCR genes in the obtained repertoire is essential to receiving the correct biological information from immune repertoire analysis.

High-throughput sequencing (HTS) of adaptive immune receptor repertoires is widely used in immunological studies (reviewed in (Minervina et al., 2019)) for the investigation of immune response to vaccines (Minervina et al., 2021; Pogorelyy et al., 2018; Sycheva et al., 2022), tumor-infiltrating lymphocytes (Gee et al., 2018; Goncharov et al., 2022; Oliveira et al., 2021), new therapeutic agents (Huang et al., 2019; Wang et al., 2018; Wilson et al., 2022), leukemia clonality and minimal residual disease monitoring (Brüggemann et al., 2019; Komkov et al., 2020; Nazarov et al., 2016; Tirtakusuma et al., 2022; Wood et al., 2018). HTS-based methods for immune repertoire profiling use either RNA or DNA as a starting material and, in most cases, use PCR for the selective enrichment of receptor sequences. DNA-based methods generally use two-sided multiplex PCR with primers annealing to multiple V- and J-genes of the rearranged receptor (Brüggemann et al., 2019; Komkov et al., 2020; Robins et al., 2009). RNA-based methods start with cDNA synthesis, usually with TCR/BCR C(constant)-genes specific oligonucleotides, followed by one-side multiplex amplification with a set of V-gene specific primers and a universal C-gene specific primer (Wang et al., 2010). Alternatively, two universal primers are used for amplification if an artificial sequence is added to the 5’ end during synthesis using a template-switch (5’-RACE) (Mamedov et al., 2013) or ligation (Oakes et al., 2017). DNA-based methods protect the repertoire from gene transcription bias and provide more comprehensive results (Barennes et al., 2020) which include most non-functional (out-of-frame) as well as functional (in-frame) rearrangements but produce high amplification bias in the course of multiplex PCR. Additionally, each T/B cell contains a single DNA copy (i.e., two target strands) of the receptor molecule in contrast to tens of single-stranded RNA copies. RNA-based methods using 5’-RACE or ligation are characterized by the lowest PCR bias as they need a single primer pair for the amplification. However, the low efficiency of adding a universal oligo to the 5’-end makes its sensitivity comparable to or even lower than DNA-based methods. The compromise between these two approaches is the RNA-based method with a one-side multiplex that has moderate amplification bias yet sufficient sensitivity (Ma et al., 2018). Most bias in one-side multiplex RNA-based approaches could be removed by using unique molecular identifiers (UMI) (Ma et al., 2018). Unfortunately, for DNA-based methods, efficient incorporation of UMIs into the initial molecule before PCR is still challenging. The only method for DNA multiplex bias correction (Carlson et al., 2013) is undirected and cost-ineffective due to the utilization of an expensive synthetic spike-in control repertoire. Here we propose an orthogonal solution for this challenge: the first fully computational algorithm for amplification bias detection and correction in adaptive immune receptor repertoires named iROAR (immune Repertoire Over Amplification Removal).

## Results

### The rationale for the Over Amplification Rate measure

Since out-of-frame TCR/BCR rearrangements do not form a functional receptor, they are not subjected to any specific clonal expansions and selection (Murugan et al., 2012). Being a passenger genomic variation, they change their initial (recombinational) clonal frequencies just randomly following the frequency changes of the second functional (in-frame) TCR/BCR allele present in the same T/B cell clone. According to the TCR/BCR loci rearrangement mechanism, the formation of in-frame and out-of-frame allele combinations in the same cell is also a stochastic and independent process in terms of V- and J-genes frequency. It leads to the conclusion that V- and J-gene frequencies among out-of-frame rearrangements must be sufficiently stable and must be equal to the initial recombination frequencies despite repertoire changes caused by various immune challenges (Fig 1). Thus, reproducible deviation of out-of-frame V- and J-gene frequencies (for the same multiplex PCR primer set) from the initial recombinational frequencies observed in the sequenced repertoire dataset is a result of artificial aberration caused by PCR amplification rather than immune repertoire evolution. Thus out-of-frame clonotypes can be considered a natural calibrator that can be used to measure amplification bias and quantitatively correct immune repertoire data.

**Figure 1.**
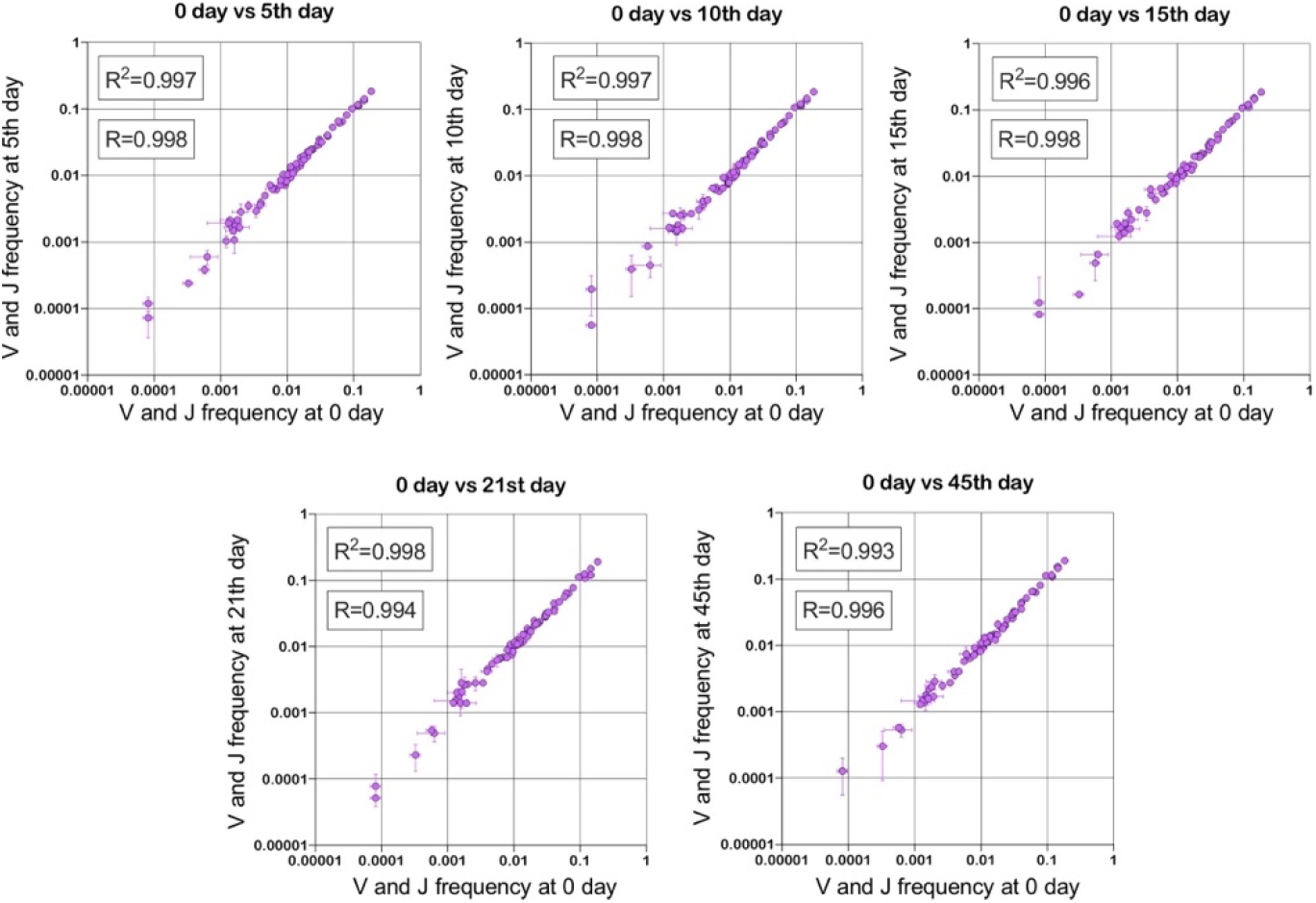
Stability of TRBV and TRBJ genes frequencies calculated based on unique out-of-frame rearrangements after Yellow fever vaccination (model of acute viral infection). Out-of-frame clonotypes for frequencies calculation were extracted from low-biased 5’ RACE TRB repertoires of PBMC samples obtained in two replicates for six time points: 0, 5, 10, 15, 21, and 45 days after YFV injection (donor M1, SRA accession number PRJNA577794 (Minervina et al., 2020)).

Formulating this observation, we developed the Over Amplification Rate (OAR) measure, which we define as a ratio of the observed and expected frequency of a V-(OAR(Vi)) or a J-gene (OAR(Ji)) among identified out-of-frame rearrangements. Observed frequency represents a value calculated as read counts (RC) for each V- and J-gene (related to out-of-frames) divided by the sum of all out-of-frame clones read count in the obtained repertoire sequencing dataset. The expected frequency is a value before amplification calculated as a number of unique out-of-frame clones (UCN) having each V- or J-gene divided by the total number of unique out-of-frame clones in the repertoire. At the final stage each OAR is normalized by dividing by the average OAR.

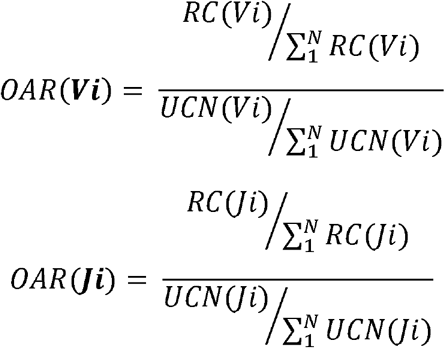

OAR value tends to be equal to 1 under ideal conditions (low or no amplification bias). It deviates from 1 as amplification bias increases in line: 5’-RACE with a single universal primer pair, one-side multiplex PCR (VMPlex), and two-side multiplex PCR (VJMPlex) (Fig 2).

**Figure 2.**
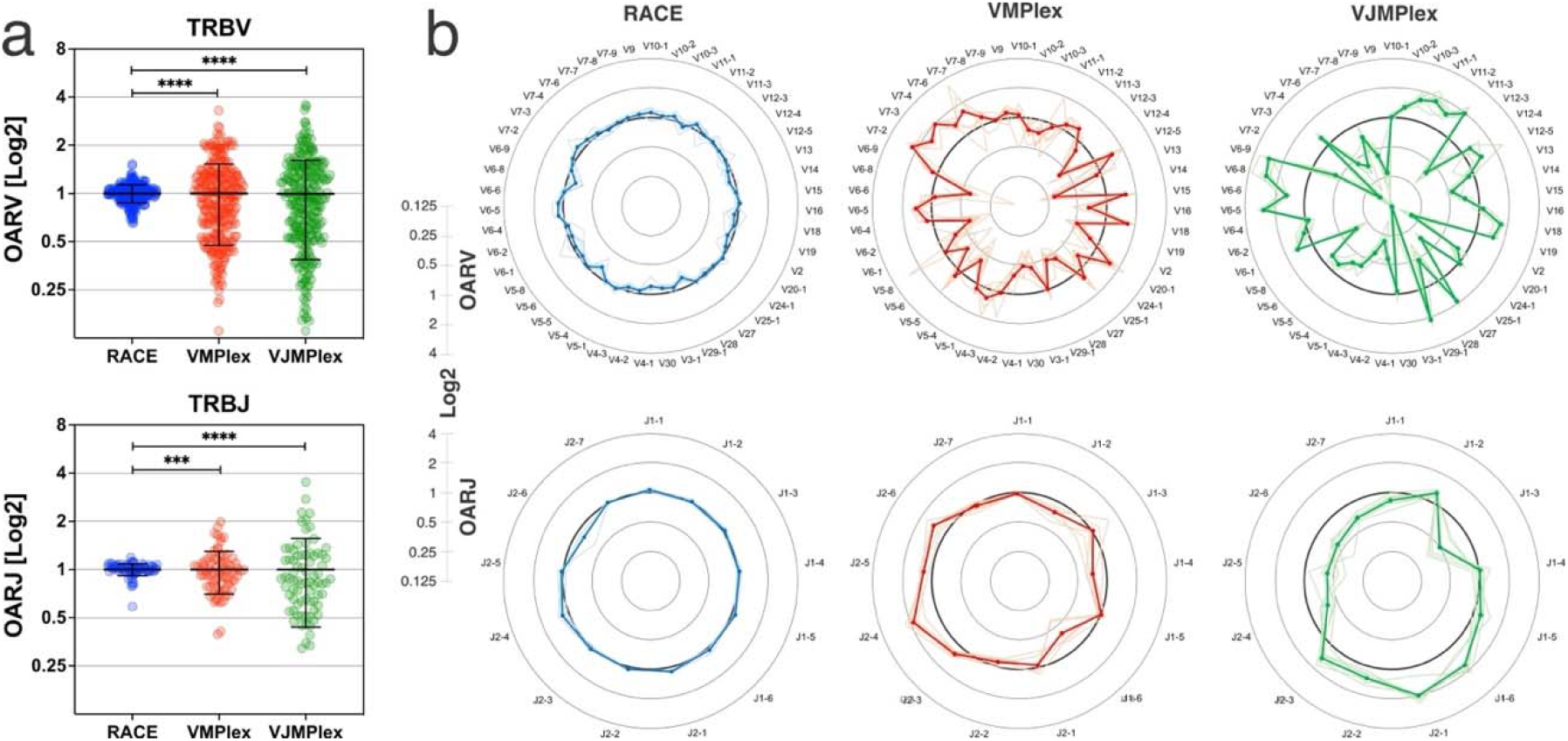
a. Comparison of OAR values variances for TRB repertoires obtained with 5’-RACE, one-side multiplex (VMPlex), and two-side multiplex (VJMPlex) PCR. The Levene’s test was performed to compare OAR variances: ****P<0.0001, ***P<0.001. The bar and whiskers indicate a mean and standard deviation. b. Average (bold lines) OAR values for TRBV and TRBJ genes in repertoires obtained with 5’-RACE, one-side multiplex), and two-side multiplex (VJMPlex) PCR. Pale lines illustrate OARs of individual repertoires. Datasets: six repertoires for RACE from PRJNA847436 (Sycheva et al., 2022), six repertoires for VMPlex from PRJNA427746 (Ma et al., 2018), six repertoires for VJMPlex from 27483#.XpCuQ1MzZQI (zenodo.org) (Weinberger et al., 2015).

### The versatility of OAR measure

OAR measurement is a universal approach and can be applied to different types of immune repertoire data. To demonstrate this versatility, we calculated OAR values for low-biased (5’ RACE) repertoires of different adaptive immune receptor chains obtained from bulk human PBMC: TCR alpha (TRA), TCR beta (TRB), and BCR heavy chains (Fig. 3a). The results show that OARs for both TCR and BCR repertoires obtained by 5’ RACE are close to 1 and stay within the range of 0.5 to 2, which is much narrower than OAR for multiplex PCR-based repertoires (see main text Fig. 2).

**Figure 3.**
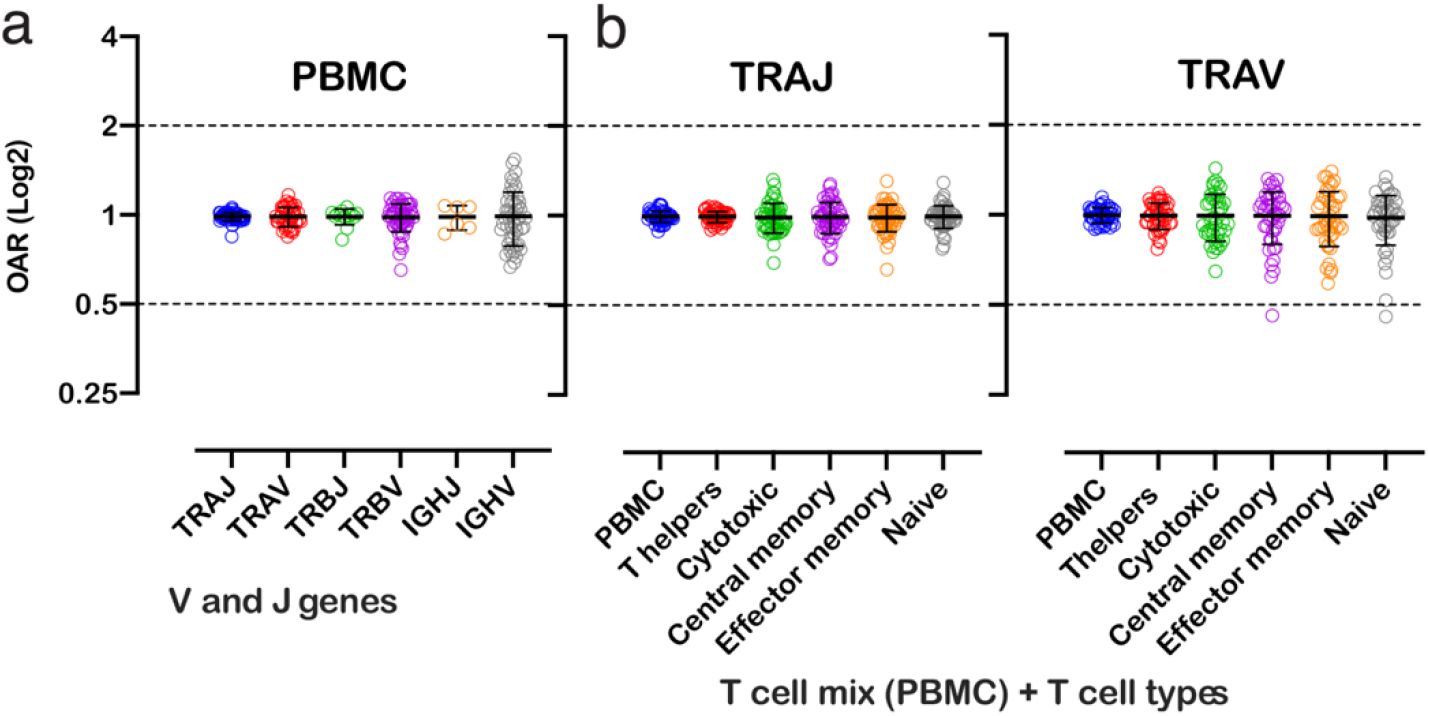
Distribution of Over Amplification Rates of V- and J-genes in RACE-based repertoires of TCR and BCR (the empty dots represent average OARs among TCR repertoires: SRA accession numbers: PRJNA577794, PRJNA316572, PRJEB27352, and BCR repertoires: SRA accession number: PRJNA297771, PRJNA494572). B. Over Amplification Rates of V- and J-genes of TCR alpha chains in RACE-based repertoires of different types of T-cells (donors M1 and P30, 45^th^ day after booster vaccination, SRA accession number PRJNA577794 (Minervina et al., 2020)) The bar and whiskers indicate a mean and standard deviation.

We also analyzed OARs for low-biased (5’ RACE) TCR repertoires of different T cell subpopulations, including T-helper, cytotoxic, central memory, effector memory, and naïve T cells. As shown in Fig. 3, the OAR values demonstrate much less differences between analyzed T cell types then between RACE and multiplex PCR and are close enough to 1 similarly to the repertoire of bulk T cell mix obtained from PBMC.

Herewith, the variance of IGHV’s OARs compared TCRs’ and the variance of TCR subpopulations OARs compared PBMCs’ is slightly higher. This phenomenon may be linked to well-known differences in clonal expansion intensities of B/T-cell subsets which can affect indirectly the OAR values. However, the proof of this hypothesis demands separate deep analysis which is beyond the main focus of this research.

Despite it, our results demonstrate that OAR is a sufficiently universal measure of repertoires and can be applied to most adaptive immune receptors and cell types.

### Factors affecting OAR measure accuracy

In the case of insufficient sequencing coverage, high PCR bias can lead to the dramatic loss of clones and thus an incorrect measurement of V- and J-genes frequencies. In this instance, for the majority of V- and J-genes, the population frequencies can approximate the real frequencies better than multiplex repertoire-based ones (Suppl Fig 1). If upon comparison samples’ UCN-based frequencies significantly differ from the average frequencies calculated for the population (i.e., exceeds 99% confidence interval), OAR calculation should be based on the latter.

Also, the balance of V- and J-genes frequencies can be disrupted by accidentally arisen abnormally large non-functional clonotypes generated in the course of abnormal clonal expansion in various lymphoproliferative disorders or stochastic spike in normal lymphocyte population. To reduce the impact of this anomaly on OAR value, the top clone of each V and J-gene containing subgroups must be excluded from OAR calculation. Since V- and J-specific bias affects all clones non-selectively, the remaining large part of clones after top clones exclusion should be still representative for PCR bias calculation. As shown in Fig4a, the exclusion of one top clonotype from OAR calculation for RACE-based TRB repertoire is enough to restore OAR calculation accuracy for TRBV2, TRBV5-6, TRBV7-9, TRBV11-3. The further top clones exclusion has no significant effect on OAR values.

Another aspect impacting the accuracy of OAR calculation is the low sequencing coverage of the TCR/BCR repertoire. The ratio of total read counts and the sum of unique clone counts can affect OAR value despite PCR bias solely because of the mathematical properties of the OAR formula. In the extreme case, the OAR value (OAR = 1) for V- and J-genes represented in a single out-of-frame clone with only one read will not reflect the real amplification bias. To address this issue, we analyzed the OAR calculation error as a function of the number of reads per clone used for the OARs evaluation (Fig. 4b). For this purpose, we performed a serial down-sampling of TCR datasets generated by RACE and two-side multiplex PCR and calculated OAR measurement error for each dataset portion. OAR calculated for the entire dataset was taken as a benchmark. The result shows that 1.8 (for MPlex) and 2.5 (for RACE) reads per out-of-frame clonotype are a minimal sufficient sequencing coverage to get adequate OAR values with an acceptable error rate of ∼10%.

**Figure 4.**
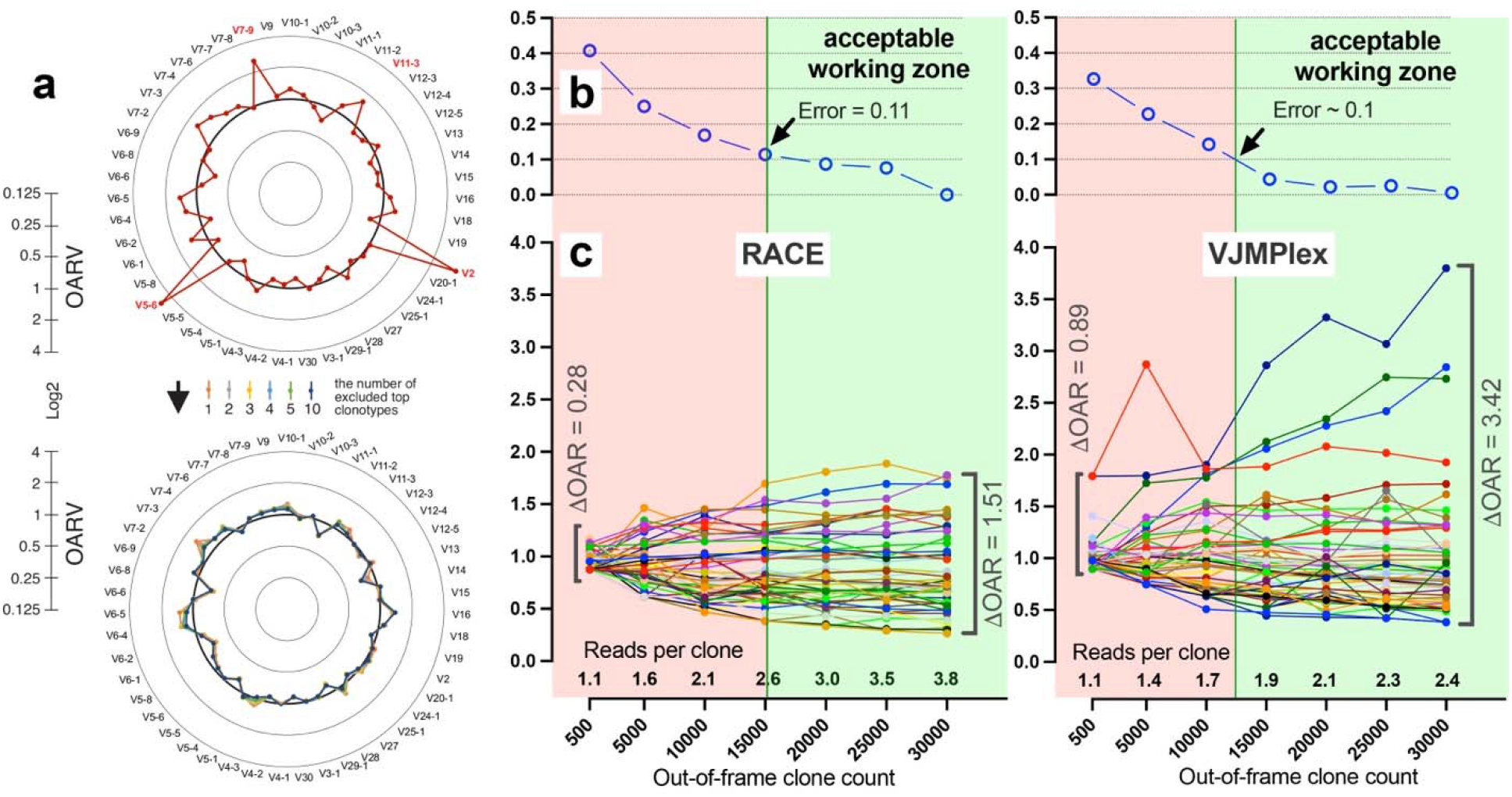
Factors impacting OAR calculation accuracy. a. Impact of highly proliferated top non-functional clonotype on OAR calculation accuracy in low-biased RACE-based TRB repertoire (Data: SRR19594184). b. Impact of sequencing depth on OARs calculation error. c. PCR bias independent changes of TRB V-genes OARs as a function of sequencing depth. Data: two-sided multiplex-based TRB repertoire (Data: RACE - SRR3129976, VJMPlex – SRR3129972).

It is also important to note that errors in nucleotide sequences occurring during library preparation and sequencing could lead to an artificial increase in both in-frame and out-of-frame clone diversity. Single nucleotide substitutions generate artificial clones as a branch of real most abundant clones inside of each in-frame and out-of-frame group independently. Single nucleotide indels lead to cross-generation artificial clones between groups: real in-frame clones generate false out-of-frame clones and vice versa. Artificial clones compromise the accuracy of both repertoire itself and OAR value. To eliminate such clones generated by single-nucleotide substitutions, we filtered them out by the VDJTOOLS software (see Methods section). To eliminate artificial clones produced by indels, we searched for in-frame and out-of-frame clone pairs which differ by one indel (Levenshtein distance = 1). If their ratio is less than 1:500, the smaller clone in pair is discarded, and its count is added to the count of the larger clone (this procedure guarantees to discard most sequencing errors present in 1 per 1000 nucleotides average).

### Over amplification rate index

To estimate the value of immune repertoire structure disruption by amplification bias, we proposed the OAR-index, which represents the mean square deviation of OARs for each V and J gene from the value characteristic for repertoire with no bias (OAR=1). OAR-index is directly proportional to the amplification bias and thus can be used for rapid estimation and comparison of immune repertoire bias. The less OAR index is, the less PCR bias is with an ideally unbiased repertoire having OAR-index = 0.

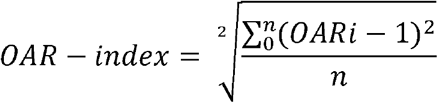

### UAR ∑ UAR 1 Using OAR for the removal of amplification bias

Normalization coefficients for each V-J combination are estimated by multiplication of corresponding V- and J-gene OARs for two-side multiplex and V-gene OAR for one-side multiplex (Fig 2a). The corrected read count for each clonotype with the particular V-J gene combination is obtained simply by dividing the observed read count by the corresponding normalization coefficient. OAR of V- and J-genes could be co-dependent, which can be a reason for overcorrection. To avoid this issue, the procedure can be recursively repeated with a modified normalization coefficient defined as described coefficient raised to the power of a number in the range from 0 to 1 (parameter “mt”). The corrected read counts are used to estimate the real percentage of each clonotype in the repertoire. However, the all multiplex-based repertoires analyzed in actual study required just 1 iteration with mt = 1. A detailed flowchart of the OAR-based amplification bias correction algorithm named iROAR is shown in Fig 5a.

**Figure 5.**
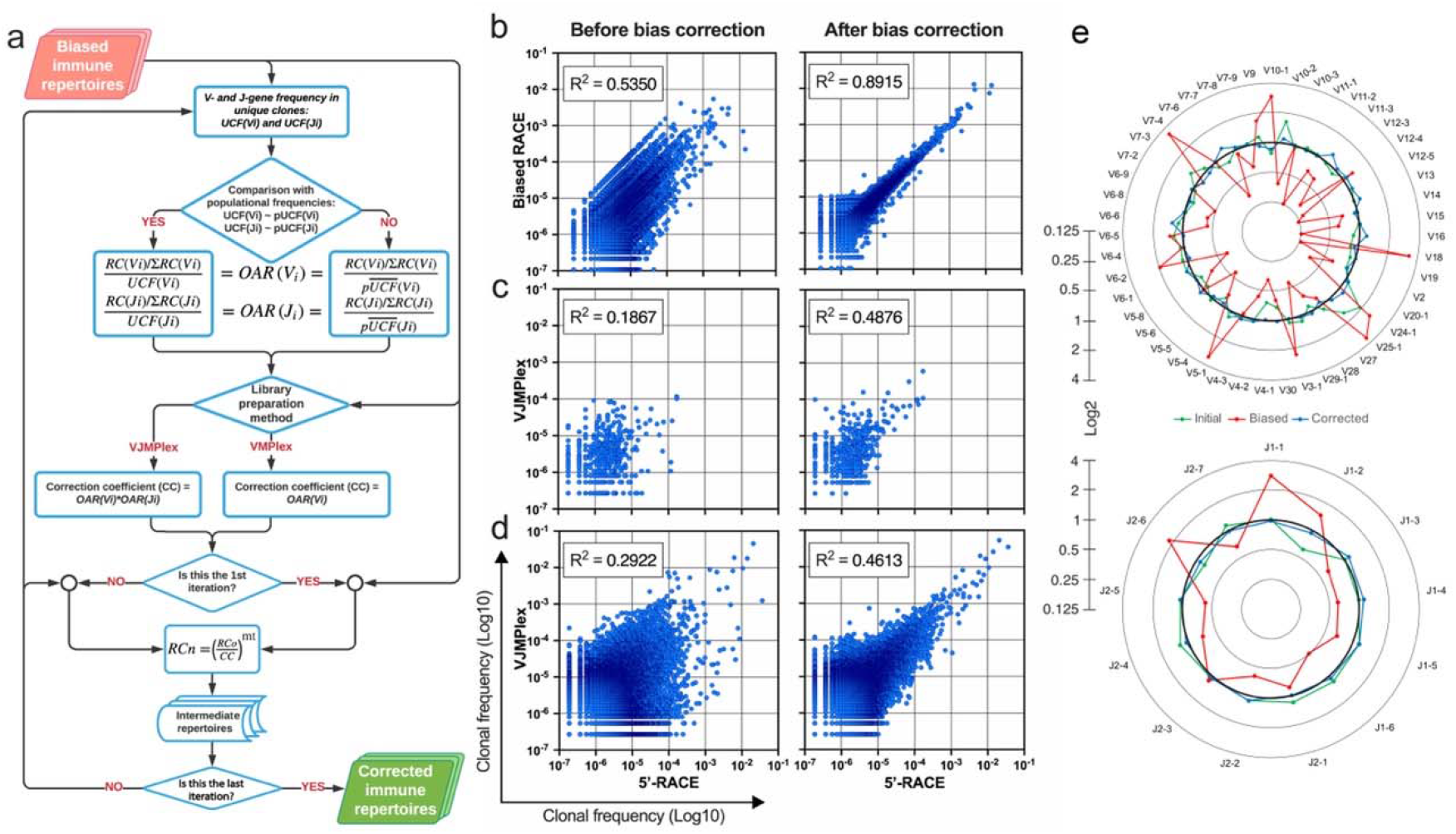
a. Flowchart of iROAR algorithm. UCF – Frequency calculated using unique clones counts (denominator of OAR), pUCF – population UCF, RC – read count, RCn – normalized RC, RCo – observed RC, mt – the number in the range from 0 to 1 for the iterative procedure. b. Clone frequencies in the low biased 5’-RACE-based repertoire (ENA database, accession number ERR2869430) versus the same repertoire with introduced artificial bias: before and after iROAR processing. c. Out-of-frame (non-functional) clone frequencies in low biased 5’-RACE-based repertoire versus two-side multiplex (VJMPlex)-based repertoire obtained for the same RNA sample (SRA database, accession numbers SRR3129976 and SRR3129972): before and after iROAR processing. d. In-frame (functional) clone frequencies in low biased 5’-RACE-based repertoire versus two-side multiplex (VJMPlex)-based repertoire obtained for the same RNA sample: before and after iROAR processing. SRA database, accession numbers SRR3129976 and SRR3129970. R^2^ is the squared Pearson correlation coefficient. iROAR was applied only for biased repertoires: artificially biased RACE and VJMPlex. e. OARV and OARJ of test 5’-RACE-based TRB repertoire (Fig.5b) before artificial bias introduction (green dots and line), biased one (red dots and line) and corrected one by iROAR (blue dots and line).

### OAR-based approach validation

The validation of OAR-based amplification bias correction was performed on the TRB dataset with *in silico* introduced bias generated from real (experimental) low-biased (5’-RACE) repertoire (Fig 5b). After correction, the OAR-index indicating general repertoire bias expectedly decreased from 1.81 to 0.76. Interestingly, the OAR independent measure -R-squared value of in silico biased and original repertoire correlation raised from 0.5350 to 0.8915 confirming the substantial reduction of *in silico* introduced quantitative bias. Afterward, we tested our approach on real paired experimental datasets obtained from the same RNA sample by two different method types: 5’-RACE and multiplex PCR (Barennes et al., 2020; Liu et al., 2016) (Fig 5c-d, Fig 6).

**Figure 6.**
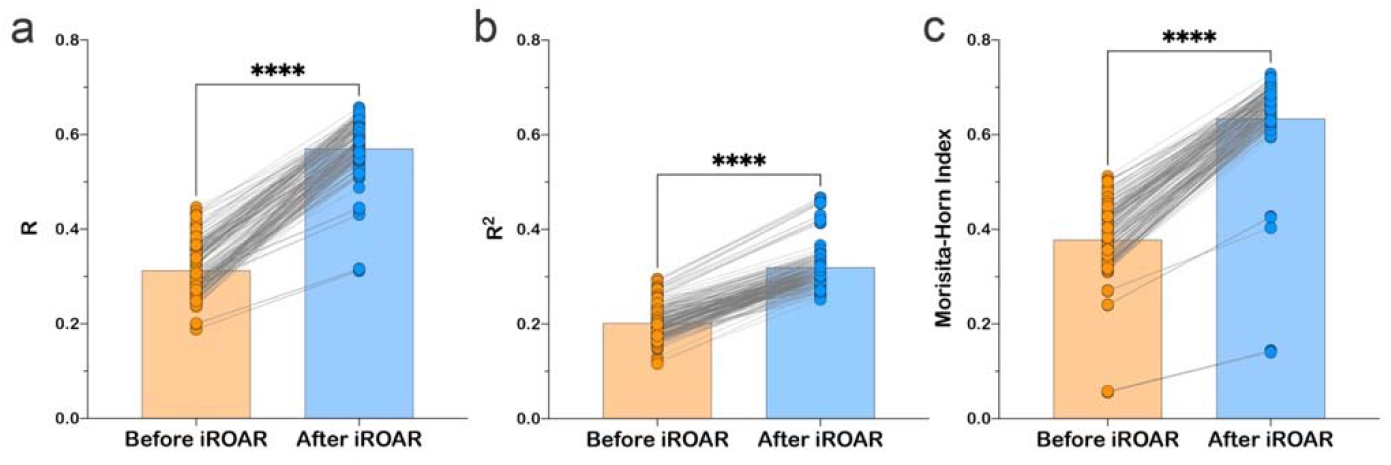
Effect of iROAR-based PCR bias correction in MPlex repertoire on similarity with low-biased RACE-based repertoire obtained from the same RNA sample. a) Pearson correlation coefficient; b) R-square measure; c) Morisita-Horn similarity index. **** P<0.0001 (two-tailed Wilcoxon matched-pairs signed rank test, CI=0.95). Dataset: PRJNA548335 (3 different RACE (RACE-2, RACE-3, RACE-4 in 6 replicates each) protocols vs. RNA-based MPlex (Multiplex-3) protocol for Donor1 and Donor2 (100 ng RNA input): total 36 points; PRJNA309577 (One RACE protocol vs. one MPlex protocol for Donors S01 (4 MPlex replicates vs. 2 RACE replicates), S02 (2 MPlex replicates vs. 4 RACE replicates) and donor S03 (1 MPlex replicate vs. 1 RACE replicate): total 17 points).

As a result of amplification bias correction, OAR index for multiplex-based repertoire decreased 1.5-fold average. At the exact time, the correlation of clonal frequencies obtained with RACE and multiplex significantly increased (Pearson correlation measure and R-squared value increased 1.5-fold average each) with a significant rise of repertoires similarity (Morisita-Horn index increased 1.7-fold average) (Fig 6). Importantly, amplification bias decreased in both out-of-frame and in-frame clone sets, although normalization coefficients ware calculated using out-of-frame ones only.

### Comparison of iROAR and spike-in-based approach for amplification bias detection

Biological spike-in is considered a classical technique for multiplex PCR bias evaluation. Several options for this technique including synthetic repertoire (Carlson et al., 2013; Wu et al., 2020), lymphoid cell lines DNA mix and DNA from human blood, tonsil, and thymus (Kallemeijn et al., 2018; Knecht et al., 2019) were established to measure V- and J-segment specific primers performance during TCR/BCR rearrangements amplification in multiplex PCR. In this study, we compared iROAR-based amplification bias evaluation with a spike-in-based approach. Similarly to ref. (Kallemeijn et al., 2018; Knecht et al., 2019) we were using natural thymic cell-derived spike-ins rather than synthetic ones. Human CD8 T-cells derived DNA was used as a target input for the libraries’ preparation. TRA rearrangements library of thymocytes were used as a source of spike-ins. Two different random mixes of TRAV and TRAJ-specific primers (0.18-4.7 μM each) were used for multiplex PCR amplification of target DNA with spike-in added. Each test library was prepared in two replicas (four test libraries total). The obtained libraries were sequenced with an average coverage of 9.88 reads per clonotype and contained 35 818 – 40 209 target and 2 298 - 3 571 spike-in clonotypes after pseudogenes removal (Table 1).

**Table 1.** The number of spike-in and target clonotypes in test TRA libraries.

| Sample | Spike-in clonotypes |  |  | Target clonotypes |  |  |
| --- | --- | --- | --- | --- | --- | --- |
|  | number | read count | coverage | number | read count | coverage |
| Primer mix 1 Replica 1 | 3 571 | 30 474 | 8.53 | 39 911 | 348 792 | 8.74 |
| Primer mix 1 Replica 2 | 2 698 | 19 420 | 7.20 | 35 818 | 303 494 | 8.47 |
| Primer mix 2 Replica 1 | 3 439 | 34 717 | 10.10 | 40 209 | 425 508 | 10.58 |
| Primer mix 2 Replica 2 | 2 298 | 24 823 | 10.80 | 33 406 | 383 615 | 11.48 |

Multiplex PCR bias of each separate V and J-gene was calculated using both iROAR and biological spike-in approaches demonstrating the high correlation level (Fig 7a and 7b) for the matched OAR/Spike-in pairs (Pearson’s r = 0.78 average) in contrast to mismatched ones (Pearson’s r = 0.46 average). VJ combination bias for both approaches was calculated by multiplying V and J-segment biases and compared using correlation analysis (Fig 7c-e). iROAR and spike-in detected VJ biases showed a strong positive correlation (Pearson’s r = 0.7182 average) for all four test TRA libraries (Fig 7c). Based on replicas comparison, the reproducibility of iROAR detected VJ bias was higher than one detected using spike-in control (Fig 7d and 7e).

**Figure 7.**
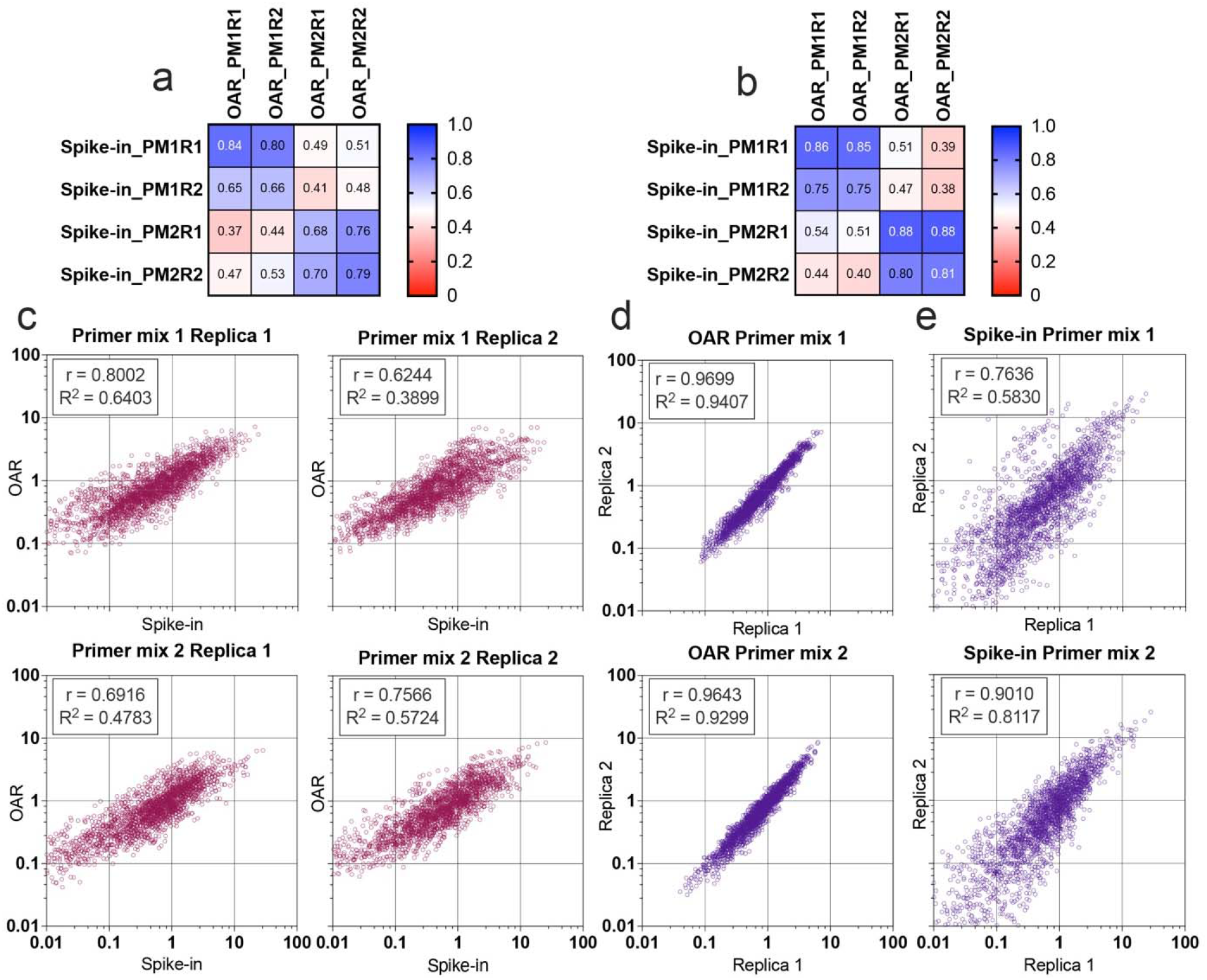
Comparison of OAR-based and biological spike-in-based approaches for multiplex PCR bias detection. Pearson’s correlation coefficient for V-segments bias measure (a) and J-segments bias measure (b). Column and row titles: PM = Primer mix, R = replica. c. Correlation of VJ combination bias calculated by iROAR and biological spike-ins. d. Reproducibility of iROAR-based VJ combination bias detection. e. Reproducibility of Spike-in-based VJ combination bias detection. Data: PRJNA825832.

### Impact of iROAR on a similarity of repertoires prepared by different multiplex PCR systems

To further test the iROAR approach’s ability to raise the uniformity of repertoires by reducing multiplex PCR-specific bias, we analyzed changes in the similarity of repertoires prepared for the same individual but using different multiplex methods. For this purpose, we compared OARs, V/J, and clonotype frequencies before and after bias correction using iROAR in test TRA libraries prepared with Primer mix 1 and Primer mix 2 (after spike-in removal). As a result of iROAR-based bias correction, the difference between OARs for these two library types significantly decreases, and OARs themselves approach a value of one. By default, iROAR does not affect the diversity of repertoires and does not remove any clonotypes. Meanwhile, V and J frequencies are subject to substantial changes (Fig.8b) depending on the initial bias level. These changes occur in both biased repertoires (Primer mix 1 and Primer mix 2) and lead to an increase its convergence (Fig.8d). Herewith R squared measure increased 1.31-fold and 1.5-fold for V and J-gene frequencies, respectively. Moreover, bias correction using iROAR also increases similarities of clone frequencies (Fig.8c). In this case both the Morisita-Horn index and Pearson correlation coefficient increase 2-fold, and R squared measure increases 1.5-fold.

**Figure 8.**
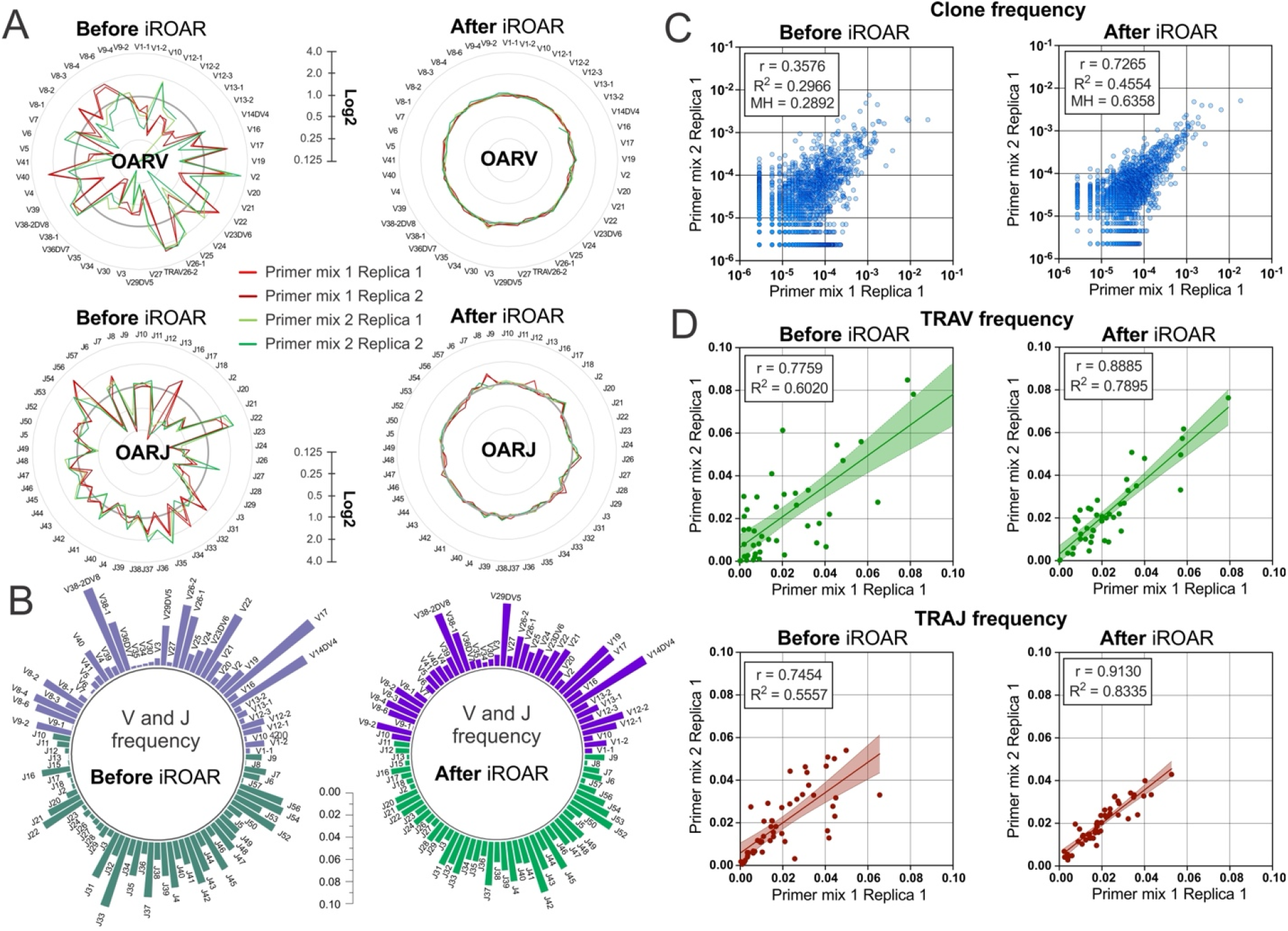
Convergence of OAR, clonotype, and V/J frequencies between two TRA repertoire before and after iROAR based bias correction. a. OAR values changes in four test TRA libraries after PCR bias correction using iROAR. b. TRAV and TRAJ frequency changes after PCR bias correction using iROAR (Sample: Primer mix 1 Replica 1). c. Correlation of clonal frequencies of two different types of test TRA repertoires before and after iROAR-based PCR bias correction d. Correlation of V and J-gene frequencies of test TRA repertoires before and after iROAR-based PCR bias correction.

It is important to note that OARs calculation and bias correction for each of the analyzed test TRA repertoires was performed entirely independently without the involvement of any common normalization coefficients or spike-in controls. Therefore, each repertoire contains enough information to correct it adequately, increasing the consistencies of interrogated repertoires obtained even by different multiplex PCR protocols.

All observed results can be considered evidence of the actual capacity of iROAR approach to accurately detect and reduce multiplex-specific quantitative bias in adaptive immune receptor repertoires.

## Discussion

Even a small difference in amplification efficiencies can lead to a massive bias after multiple amplification cycles due to the exponential nature of PCR. Thus, most of the existing immune repertoire library preparation methods are subjected to amplification bias. The effect of distinct PCR bias-generating factors can be reduced experimentally by varying reaction mixture content and introducing special protocols (UMI, crafty primer structures, spike-in controls). However, the criteria for estimating and removing the residual bias after applying these optimization approaches are lacking. Here we close this gap by introducing the OAR value and OAR-index, which score PCR bias for both V- and J-genes separately (OAR values) and the whole repertoire dataset (OAR-index). Based on OAR values, we developed the first fully computational approach to decipher and correct amplification bias in adaptive immune receptor repertoires produced by one-side or two-side multiplex PCR-based methods, using RNA or DNA as a template. Due to the inability to use UMI-based correction for DNA-based multiplex, the developed approach is the only currently available technique allowing direct measuring and correcting PCR bias in such repertoires without additional experiments.

In contrast to cell-line mix spike-in (Knecht et al., 2019) or synthetic repertoire-based (Carlson et al., 2013; Wu et al., 2020) PCR bias correction, the proposed approach operates with hundreds and even thousands of natural calibrators (out-of-frame clones) for each V-J gene pair. It makes this method potentially more reliable due to the ability to minimize the impact of CDR3 structure on PCR bias calculation since out-of-frame captures significantly higher CDR3 diversity than biological spike-ins. Moreover, similarly to a previously described method (Carlson et al., 2013) the OAR-based approach can also be used for primer efficacy evaluation to optimize their structures and concentrations, which in turn will straighten the coverage of various V- and J-genes and minimize the number of experimentally lost clones. Being fully computational, the developed PCR bias correction algorithm can be easily implemented in any TCR/BCR repertoire analysis pipeline, noticeably improving the quantitative parts of the analysis. Even though it’s not possible to fully substitute the low-biased RACE methods, iROAR is capable to make multiplex-PCR-based repertoires more consistent with RACE-based ones. Therefore, the developed approach can provide the opportunity to compare the immune repertoire datasets generated using different library preparation methods.

## Methods

### Raw-data processing and immune repertoires reconstruction

All sequencing data used in this study represent human TCR and BCR repertoires. The repertoires (see Suppl. Table 1) were reconstructed from fastq data using MiXCR v2 software (Bolotin et al., 2017, 2015) after primers and adapters trimming using FASTP software (Chen et al., 2018). All obtained repertoires were converted to VDJTOOLS (Shugay et al., 2015) format for unification. Erroneous clones generated by single nucleotide substitution were removed from the repertoire using the “Correct” function from the VDJTOOLS software package. Erroneous clones generated by single nucleotide indels were removed from repertoires using the “Filter” function from developed iROAR software. V and J pseudogenes were removed from repertoires using the “FilterBySegment” function of VDJTOOLS.

### TRA repertoires preparation

The peripheral blood was collected from a healthy volunteer from the article’s co-authors with informed consent in a certified clinical lab. PBMC was separated from whole blood using the Ficoll-Paque approach. CD8+ T-cells were isolated using Dynabeads™ CD8 Positive Isolation Kit (Invitrogen). DNA for library preparation was extracted from CD8+ T-cells using FlexiGene DNA Kit (Qiagen). 150 ng aliquots of obtained DNA were used as input to prepare each out of four TRA libraries. Each DNA aliquot was premixed with 0.1 pg of serial diluted low-biased TRA library (prepared using MiLaboratories Human TCR kit) of thymic cells (spike-in matrix) as biological spike-ins. Two pools of previously designed (Komkov et al., 2020) TRAV and TRAJ-specific primers (MiLaboratories LLC) with randomly selected concentrations (0.18-4.7 μM each) were generated to produce two types of TRA libraries with different quantitative bias status simulating libraries produced by different multiplex PCR methods. Library preparation was performed according to the protocol from ref. (Komkov et al., 2020). Both types of TRA libraries were prepared in two replicas and sequenced along with a spike-in matrix library on the MiSeq Illumina instrument (SE 150 nt) with moderate coverage 480 000 reads per library.

### Biological spike-in detection and analysis

TRA repertoires were extracted from FASTQ files using MIXCR software. All obtained MIXCR output files were converted to VDJTOOLS format as described above. Extraction of the spike-in sequences and spike-in free repertoires from sequenced libraries was performed using the VDJTOOLS function “ApplySampleAsFilter” and the sequenced spike-in library as a filter. Spike-in-based amplification bias was calculated as the quotient of V and J-frequency in spikeins extracted from target libraries and corresponding V and J-frequency in the spike-in matrix, which was not subjected to multiplex amplification. OARs for obtained TRA libraries were calculated using iROAR software and spike-in-free repertoires as input. VJ bias values were calculated by multiplying V-to J-segment-specific biases. Correlation analysis of iROAR and Spike-in VJ bias values was performed using GraphPad Prism9 software.

### Step-by-step pipeline for the OAR evaluation used in this study

1. Single nucleotide error correction in read1/read2 intersected sequences and Illumina adapters removal (optional): fastp -c -i input_R1.fastq.gz -I input_R2.fastq.gz -o fp.input_R1.fastq.gz -O fp.input_R2.fastq.gz
2. Raw reads alignment (essential):
  a. For TCR beta chains mixcr align -c TRB fp.input_R1.fastq.gz fp.input_R2.fastq.gz output1.vdjca
  b. For TCR beta chains mixcr align -c TRA fp.input_R1.fastq.gz fp.input_R2.fastq.gz output1.vdjca
3. Clonotypes assemble (essential): mixcr assemble output1.vdjca output2.clns
4. TCR repertoire export in a human-readable format (essential): mixcr exportClones output2.clns clones.txt
5. Convert repertoire into VDJtools format (essential): java -jar vdjtools.jar Convert -S mixcr clones.txt vdjtools
6. Artificial diversity removal by single nucleotide substitutions correction (optional): java -jar vdjtools.jar Correct vdjtools.clones.txt correct
7. Pseudogenes removal (optional):
  a. For TCR beta chains java -jar vdjtools.jar FilterBySegment –-j-segment TRBJ2-2P --v-segment TRBV1,TRBV12-1,TRBV12-2,TRBV17,TRBV21-1,TRBV22-1,TRBV23-1,TRBV26,TRBV5-2,TRBV5-3,TRBV5-7,TRBV6-7,TRBV7-1,TRBV7-5,TRBV8-1,TRBV8-2 –negative correct.vdjtools.clones.txt filter
  b. For TCR alpha chains java -jar vdjtools.jar FilterBySegment -–j-segment TRAJ1,TRAJ19,TRAJ2,TRAJ25,TRAJ51,TRAJ55,TRAJ58,TRAJ59,TRAJ60,TRAJ61 --v-segment TRAV11,TRAV11-1,TRAV14-1,TRAV15,TRAV28,TRAV31,TRAV32,TRAV33,TRAV37,TRAV46,TRAV8-5,TRAV8-6- 1,TRAV8-7 --negative correct.vdjtools.clones.txt filter
8. Artificial diversity removal by single nucleotide indels correction (optional): iroar Filter -se 0.01 filter.correct.vdjtools.clones.txt filter2.txt
9. OARs calculation and quantitative bias correction (essential): iroar Count -min_outframe 15 -r -z 1 -iter 1 -mt 1 input_folder output_folder *(input_folder must contain filter2*.*txt file)*

### OAR evaluation and statistical analysis

For the OAR and OAR-index calculation and amplification bias removal, we used the command-line-based iROAR software designed in this study and freely available for nonprofit use at GitHub (https://github.com/smiranast/iROAR). For the OAR comparison between 5’-RACE, one-side, and two-side multiplex PCRs, an equal number of out-of-frame clones (50,000) was randomly selected from TCR repertoires of 15 healthy individuals (for each approach). Average population V- and J-gene frequencies (unweighted) were calculated based on out-of-frame clones from 105 TRB repertoires obtained by two methods: 5’-RACE (95 repertoires) and single-cell TCR profiling (10X genomics) (10 repertoires) (Suppl. Table 1) using the “CalcSegmentUsage” function with “-u” parameter of VDJTOOLS. All statistical tests were performed using Prism9 GraphPad software (https://www.graphpad.com/).

### iROAR software requirement

Recommended system configuration for iROAR running: Linux or MacOS, 2 CPU, 8GB RAM, programming language: python=3.7.3, required Python packages: matplotlib=3.0.3, numpy=1.16.2, pandas=0.24.2, requests=2.21.0. Starting iROAR package includes list of average populational frequencies with standard deviations of TRB and TRA V- and J-genes related to the European population. iROAR run command: iroar Count [optional parameters] <input> <output>. Recommended parameters for most tasks: -min_outframe 15 -r -z 1 -iter 1 –mt 1. Full list of available parameters is deposited in project directory at github (https://github.com/smiranast/iROAR)

## Data access

All analyzed datasets were downloaded from open-source databases: NCBI SRA (https://www.ncbi.nlm.nih.gov/sra), ENA (https://www.ebi.ac.uk/ena), and Zenodo project (https://zenodo.org/). A complete list of web links and accession numbers is summarized in Suppl. Table 1. TRA repertoire dataset generated in this study for iROAR validation is available under access number PRJNA825832.

The iROAR software and its documentation are available at the link: https://github.com/smiranast/iROAR. The additional software used in this study is available in the GitHub repository (https://github.com/milaboratory/mixcr, https://github.com/mikessh/vdjtools, https://github.com/OpenGene/fastp)

## Acknowledgments

We thank Grigory Armeev and Valery Novoseletsky for their help with data storage and processing.

## Supplementary Data

**Supplementary Table 1**. XLSX. The file contains accession numbers and links to the datasets used in this study.

**Supplementary Table 2**. XLSX. Comparison of iROAR algorithm with the existing approaches for PCR bias removal in human adaptive immune receptor repertoires

**Supplementary Figure 1**. JPEG. V- and J-gene frequencies among unique out-of-frame rearrangements calculated using biased (two-side multiplex, empty circles) and low-biased (RACE, filled circles) are compared to average population frequencies (violin plot). Significantly biased V and J genes are highlighted pink. Paired (the same starting RNA samples) two-side multiplex and RACE data: SRA accession number PRJNA309577, average population frequencies were calculated using a series of RACE TCR and single-cell TCRseq data (see Suppl. Table 1).

