## Supplementary figures and images for "The use of non-functional clonotypes as a natural calibrator for quantitative bias correction in adaptive immune receptor repertoire profiling"

### Supplementary Figure 1

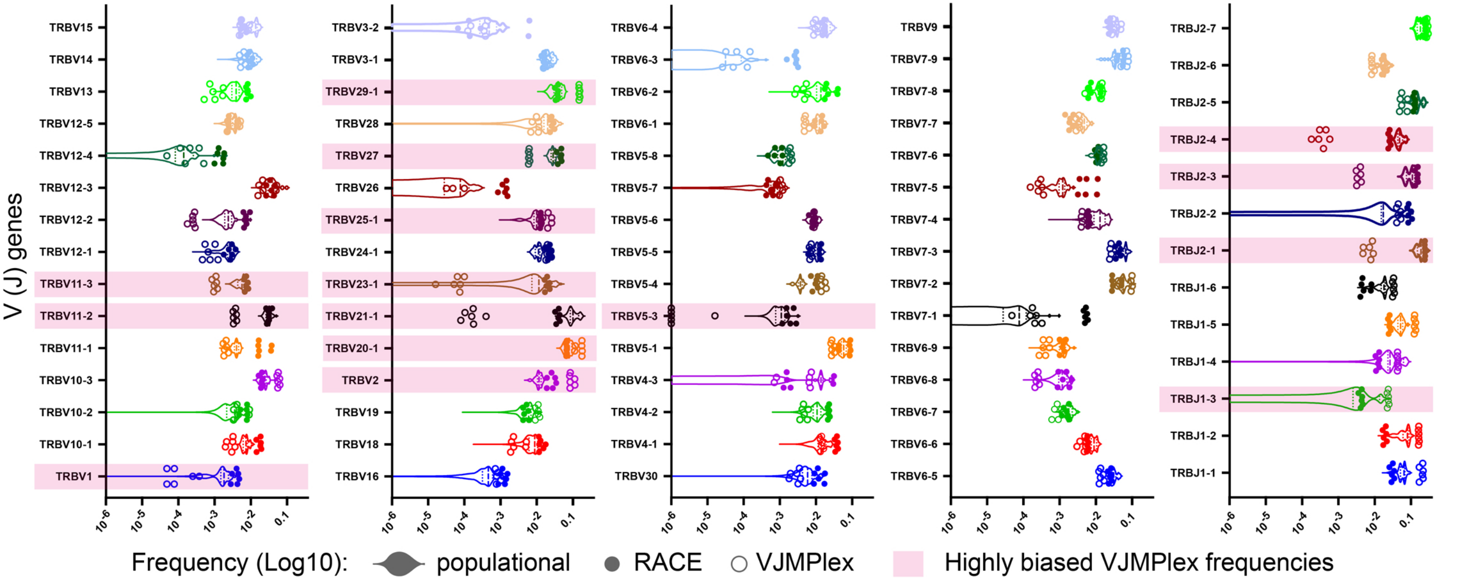
