## Supplementary Table 1 for "The use of non-functional clonotypes as a natural calibrator for quantitative bias correction in adaptive immune receptor repertoire profiling"

| Referense | PMID | SRA data accession number or link | Method |
|---|---|---|---|
| Sycheva et al., Front Immunol., 2022 | 36091004 | PRJNA847436 | 5'-RACE |
| Turchaninova et al., Nat Protocol., 2016 | 27490633 | PRJNA297771 | 5'-RACE |
| Minervina et al., eLife, 2020 | 32081129 | PRJNA577794 | 5'-RACE |
| Simon et al., J Immunol Methods., 2018 | 30312601 | PRJNA494572 | 5'-RACE |
| Ma et al., Front Immun., 2018 | 29467754 | PRJNA427746 | VMPlex |
| Weinberger et al., PLoS ONE, 2015 | 26600245 | https://zenodo.org/record/27483#.XpCuQ1MzZQI | VJMPlex |
| Liu et al., PLoS ONE., 2016 | 27019362 | PRJNA309577 | VJMPlex + 5'-RACE |
| Barennes et al., Nat Biotech., 2020 | 32895550 | PRJNA548335 | VJMplex + 5'-RACE |

| Cell type |
|---|
| T cells |
| B cells |
| T cells |
| B cells |
| T cells |
| T cells |
| T cells |
| T cells |
