## Supplementary Table 2 for "The use of non-functional clonotypes as a natural calibrator for quantitative bias correction in adaptive immune receptor repertoire profiling"

**Comparison of iROAR algorithm with the existing approaches for PCR bias removal in human immune receptor repertoires**

| | UMI based approach | Synthetic repertoire-based approach | iROAR |
|---|---|---|---|
| Cost effective | Yes | No | Yes |
| Fully computational | No* | No** | Yes |
| Applicable for RNA-based repertoires | Yes | Yes | Yes |
| Applicable for DNA-based repertoires | No*** | Yes | Yes |
| Applicable for VMplex | Yes | Yes | Yes |
| Applicable for VJMplex | No*** | Yes | Yes |
| Primer bias removal | Yes | Yes | Yes |
| Primer efficacy evaluation | Yes | Yes | Yes |
| Direct method | Yes | No** | Yes |

* Requires specific experiment design to attach UMI to molecules before PCR

** Requires upfront experiments with synthetic repertoires

*** Are not currently developed
